# Swallowing and chewing difficulties are associated with presynaptic dopaminergic dysfunction and greater non-motor symptom burden in early drug-naïve Parkinson’s patients

**DOI:** 10.1101/577148

**Authors:** Sotirios Polychronis, Georgios Dervenoulas, Tayyabah Yousaf, Flavia Niccolini, Gennaro Pagano, Marios Politis

**Author notes:** Correspondence & reprint requests* to Marios Politis, MD, MSc, PhD, Neurodegeneration Imaging Group (NIG), Department of Clinical Neuroscience, Maurice Wohl Clinical Neuroscience Institute, Institute of Psychiatry, Psychology and Neuroscience (IoPPN), King’s College London, 125 Coldharbour Lane, Camberwell, London SE5 9NU, UK.

## Abstract

**Background:** The underlying pathophysiology of swallowing and chewing difficulties is multifactorial and evidence clarifying the precise mechanisms are scarce. Dysfunction in dopamine-related and non-dopamine-related pathways, changes in cortical networks related with swallowing and peripheral mechanisms have been implicated in the pathogenesis of swallowing difficulties. We aimed at investigating whether swallowing and chewing difficulties are associated with presynaptic dopaminergic deficits, faster motor symptom progression and cognitive decline in a population of early drug-naïve patients with Parkinson’s disease.

**Methods:** By exploring the database of Parkinson’s Progression Markers Initiative we identified forty-nine early drug-naïve Parkinson’s disease patients with swallowing difficulties. Swallowing and chewing impairment was identified with SCOPA-AUT question 1 (answer regularly) and was assessed with MDS-UPDRS Part-II, Item 2.3 (Chewing and Swallowing). We compared Parkinson’s disease patients with swallowing and chewing difficulties to Parkinson’s disease patients without difficulties, and investigated differences in striatal [^123^I]FP-CIT single photon emission computed tomography levels. Using Cox proportional hazards analyses, we also evaluated whether swallowing impairment can predict motor deterioration and cognitive dysfunction.

**Results:** Patients with Parkinson’s disease, harbored a greater deterioration regarding motor and non-motor symptoms and decreased [^123^I]FP-CIT binding when compared with patients without swallowing and chewing impairment. Higher burden of swallowing and chewing dysfunction (MDS-UPDRS-II, item 2.3) was correlated with lower [^123^I]FP-CIT uptakes within the striatum (r_s_=−0.157; *P=*0.002) and the caudate (*r*_*s*_=−0.156; P=0.002). The presence of swallowing and chewing difficulties was not a predictor of motor progression (Hazard ratio [HR]: 1.143, 95% confidence interval [CI]: 0.848–1.541; *P*=0.379) or cognitive decline (HR: 1.294, 95% CI: 0.616–2.719; *P*=0.496).

**Conclusions:** Swallowing and chewing impairment is associated with decreased presynaptic dopaminergic integrity within caudate and greater motor and non-motor symptoms burden in early drug-naïve PD.

**Author contributions:** S.P. and M.P. conceived the study, conceptualized the experimental design. M.P., S.P., G.D. and G.P. gave input to experimental design. S.P. wrote the first draft and prepared the manuscript. G.P. and S.P. performed the statistical analysis. G.P., G.D., T.Y. and S.P. generated the figures. F.N., M.P., S.P., G.P., G.D. interpreted the data. All authors revised and gave input to the manuscript.

**Financial Disclosure Statement:** Data used in the preparation of this article were obtained from the Parkinson’s Progression Markers Initiative (PPMI) database (www.ppmi-info.org/data). For up-to-date information on the study, visit www.ppmi-info.org. PPMI – a public-private partnership - is sponsored by the Michael J. Fox Foundation for Parkinson’s Research (MJFF) and is co-funded by MJFF, Abbvie, Avid Radiopharmaceuticals, Biogen Idec, Bristol-Myers Squibb, Covance, Eli Lilly & Co., F. Hoffman-La Roche, Ltd., GE Healthcare, Genentech, GlaxoSmithKline, Lundbeck, Merck, MesoScale, Piramal, Pfizer and UCB.PPMI. Industry partners are contributing to PPMI through financial and in-kind donations and are playing a lead role in providing feedback on study parameters through the Industry Scientific Advisory Board (ISAB). Through close interaction with the study, the ISAB is positioned to inform the selection and review of potential progression markers that could be used in clinical testing.

Mr. Polychronis, Dr. Dervenoulas, Ms Yousaf, Dr. Niccolini, Dr. Pagano and Prof. Politis report no disclosures.

**Potential Conflicts of Interest:** No potential conflict of interest relevant to this article was reported.

## INTRODUCTION

Swallowing and chewing difficulties are characterised by the inability to safely swallow fluids and /or solid food and are accounting for severe complications, commonly aspiration pneumonia that substantially increases mortality rates. Swallowing and chewing difficulties are more likely to have a neurologic basis and are an important issue in Parkinson’s disease (PD).(1) Between 70 to 100% of patients with PD have swallowing and chewing difficulties throughout the disease progression. The prevalence increases proportionally in later stages of the disease.(2–4) Aspiration pneumonia in the context of swallowing impairment is regarded one of the most important factors contributing to decreased life expectancy of PD patients. In addition, malnutrition, dehydration and medication intake complications are further consequences of swallowing impairment that contribute to significant decline in the quality of patients life.(5) The pathophysiology underlying swallowing and chewing difficulties in PD remains elusive. Several cortical, subcortical and peripheral mechanisms have been partially implicated without a robust and conclusive justification. The swallowing process involves multiple cortical areas, including primary sensorimotor cortex, sensorimotor integration areas, insula, anterior cingulate cortex and the supplementary motor cortex.(6–9) Swallowing and chewing impairment can also be attributed to brainstem pathology. Notably, early in the course of PD, areas that generate the central swallowing pathway in the medulla (motor nucleus of glossopharyngeal nerve, vagus nerve, reticular activating system) are exposed to the neurodegenerative processes. (10, 11) In addition, the pedunculopontine tegmental nucleus, receives abnormal inhibitory through the pallidum and is thus further exposed to neurodegeneration. (12)

In this study we investigated the association of swallowing and chewing impairment and dopaminergic deficits using [^123^I]FP-CIT single photon emission computed tomography (SPECT). Finally, we explore whether swallowing and chewing difficulties were a predictor of motor symptom progression and cognitive decline.

## METHODS

### Subjects and clinical evaluation

From the 412 PD patients included in the Parkinson’s Progression Markers Initiative (PPMI) database (www.ppmi-info.org/data), a total of 398 early drug-naïve PD patients underwent both [^123^I]FP-CIT SPECT assessments, and therefore were integrated in our analytical approach. Among these 398 PD patients, we identified 307 cognitively intact (MoCA≥26), with a complete 60-month follow-up and we included them for the longitudinal analysis. All PD patients were recruited between 2010-2015, diagnosed with PD less than two years prior to a screening visit, never treated with dopamine replacement therapy and presented with two among bradykinesia, resting tremor and rigidity or with asymmetric resting tremor/bradykinesia at screening. The diagnosis was confirmed by the presence of dopaminergic deficit at [^123^I]FP-CIT SPECT imaging.

The presence of swallowing and chewing difficulties was identified with the SCOPA-AUT question 1 (In the past month, have you had difficulty swallowing or have you choked? Answer: regularly) and quantified according to the Movement Disorder Society-sponsored revision of the Unified Parkinson’s Disease Rating Scale (MDS-UPDRS) Part-II, Item 2.3 (Chewing and Swallowing) ≥ 1. This item is a clinician-based scale consisting of 5 scores, rating between 0 (normal) and 4 (most severe impairment). Motor symptom burden was measured with the MDS-UPDRS-III and staged with the Hoehn and Yahr (H&Y) scale. Each motor domain (bradykinesia, resting tremor, rigidity, postural instability) was calculated using specific MDS-UPDRS-III sub-items as follows: bradykinesia (Total score range 0–52) = sum of Item 3.4 finger tapping, item 3.5 hand movements, item 3.6 pronation-supination movements of hands, item 3.7 toe tapping, item 3.8 leg agility, item 3.9 arising from chair, item 3.13 posture and item 3.14 body bradykinesia; rigidity (Total score range 0–20) = sum of Item 3.3 rigidity (neck, upper limbs and lower limbs); resting tremor (total score range 0–24) = sum of item 3.17 rest tremor amplitude (lip/jaw, upper limbs and lower limbs) and item 3.18 constancy of tremor; axial (total score range 0–12) = sum of item 3.10 gait, item 3.11 freezing of gait and item 3.12 postural stability (PPMI, 2011). MDS-UPDRS-II score was calculated excluding Item 2.3 (Chewing and Swallowing).

PD motor phenotypes were identified as either tremor-dominant or akinetic-rigid by applying a numerical ratio derived from the mean score of tremor and the mean score of rigidity-akinesia. (13) Patients with ratio < 0.8 were classified as akinetic-rigidity phenotype, patients with ratio > 1.0 were classified as tremor-dominant phenotype and patients with ratio between 0.8 and 1 were classified as mixed subtype. Non-motor symptoms were assessed using MDS-UPDRS-I and the Scale for Outcomes for PD– Autonomic function (SCOPA-AUT). Neuropsychiatric symptoms were assessed with the short version of the 15-item Geriatric Depression Scale (GDS) and the State Trait Anxiety Total scale (STAI). Sleep disorders were assessed with the Epworth Sleeping Scale and REM sleep behavior disorder questionnaire (RBDQ). Cognitive impairment was assessed with the Montreal cognitive assessment (MoCA). Olfactory dysfunction was assessed with the University of Pennsylvania Smell Identification Test (UPSIT). Disability was estimated using the Modified Schwab & England Activity of Daily Living (ADL). An annual assessment of cognition included scales exploring four major cognitive domains including memory [Hopkins Verbal Learning Test-Revised (HVLT-R) Recall, HVLT-R Recognition Discrimination], visuospatial functions [Judgment of Line Orientation (Benton)], working memory and executive functions [Letter Number Sequencing (LNS), Semantic Fluency], attention and processing speed [Symbol Digit Modalities Test (SDMT)].

### Dopaminergic imaging

SPECT images were obtained 4±0.5 hours after administrating an injection of approximately 185 MBq [^123^I]FP-CIT. [^123^I]FP-CIT SPECT scans were analysed following the imaging technical operations manual (http://ppmi-info.org/). Raw SPECT data was acquired into a 128 x 128 matrix stepping each 3 degrees for a total of 120 (or 4 degrees for a total of 90) projections in a window centred on 159±10%KeV. The total scan duration was 30-45 minutes. A Chang 0 attenuation correction was applied using a customised *Mu* determined empirically from the anthropomorphic brain phantom acquired at each site. A standard Gaussian 3D 6.0mm filter was applied to each image volume and then normalised to standard Montreal Neurologic Institute space. Each scan was interpreted by two independent readers who were blinded to the subjects’ demographics and characteristics. For quantification, SPECT image volumes were spatially normalized to an Ioflupane template. The eight most prominent axial slices containing striatum were summed and then a standardized volume of interest (VOI) template was applied to this image. VOI analyses were performed on the left and right caudate and putamen with the occipital region serving as a reference tissue. Specific binding ratios (SBR) were calculated as the ratio of the caudate or putamen VOI count density divided by count density of the occipital cortex minus 1. This measure approximates the binding potential, BP_ND_, when the tracer is in equilibrium at the target site and was previously reported with Ioflupane SPECT.(14)

### Assessment of motor progression and cognitive decline

Motor progression was defined as a change of one point in the H&Y scale at the follow-up visits. Cognitive decline was defined as having a clinical deterioration of cognitive function reported by the patient or the caregiver, a MoCA score<26 and at least 2 test scores (of six neuropsychological tests indicated above; regardless of the domain tested). Scores should be above 1.5 standard deviation and below the standardized mean scores of education and age. Norms were applied according to current literature (15). Follow-up visits took place in the outpatient unit of the reference hospitals once every 6 months and 307 early drug-naïve PD patients were followed up for an average of 60 months.

### Standard protocol approvals, registrations, and patient consents

This study is registered with ClinicalTrials.gov (No: NCT01141023). Each PPMI site has received approval from an ethical committee on human experimentation before the study’s initiation. Written informed consent for research was obtained from all individuals participating in the study. The present study was performed according to the STROBE guidelines.(15)

### Statistical analysis

Statistical analysis and graph illustration were performed with SPSS (version 20) and GraphPad Prism (version 6.0c) for MAC OS X, respectively. For all variables, variance homogeneity and Gaussianity were tested with Kolmogorov-Smirnov test.

Multivariate analysis of variance (MANOVA) was used to assess the main differences in clinical, imaging and non-imaging parameters between PD patients with and without swallowing and chewing difficulties. If the overall multivariate test was significant, *P*-values for each variable were calculated following Bonferroni’s multiple comparisons test. Categorical variables are expressed as proportions and compared using the χ2 test. We interrogated correlations between the swallowing and chewing scores and imaging data using Spearman’s rank correlation. To explore whether swallowing and chewing impairment can predict motor disease burden and cognitive dysfunction, Cox proportional hazards analyses were carried out. The time to occurrence of the first event for a given subject was used in the Cox model. All data are presented as mean ± standard deviation (SD), and the level α was set for all comparisons at *P*<0.05, corrected.

## RESULTS

### Clinical characteristics

The prevalence of swallowing and chewing difficulties in the cohort of early drug-naïve PD patients was 12.3% (49/398). No significant differences were observed for demographic characteristics (age: *P*>0.1, gender: *P*>0.1, disease duration: *P*>0.1 and family history of PD: *P*>0.1) between the two groups.

Swallowing and chewing difficulties were more common in early drug-naïve PD patients with akinetic-rigid motor phenotype (34/49; 69.4% *vs* 207/349; 59.3%). However, no significant difference was observed in H&Y stage (*P*>0.1), MDS-UPDRS-III (*P*>0.1), bradykinesia (*P*>0.1), rigidity (*P*>0.1), axial (*P*>0.1) and resting tremor (*P*>0.1) subscores between early drug-naïve PD patients with and without swallowing and chewing difficulties (Table 1). Early drug-naïve PD patients with swallowing and chewing difficulties had significant higher burden in motor aspects of daily living as measured by the MDS-UPDRS-II (*P*<0.001).

**Table 1.**
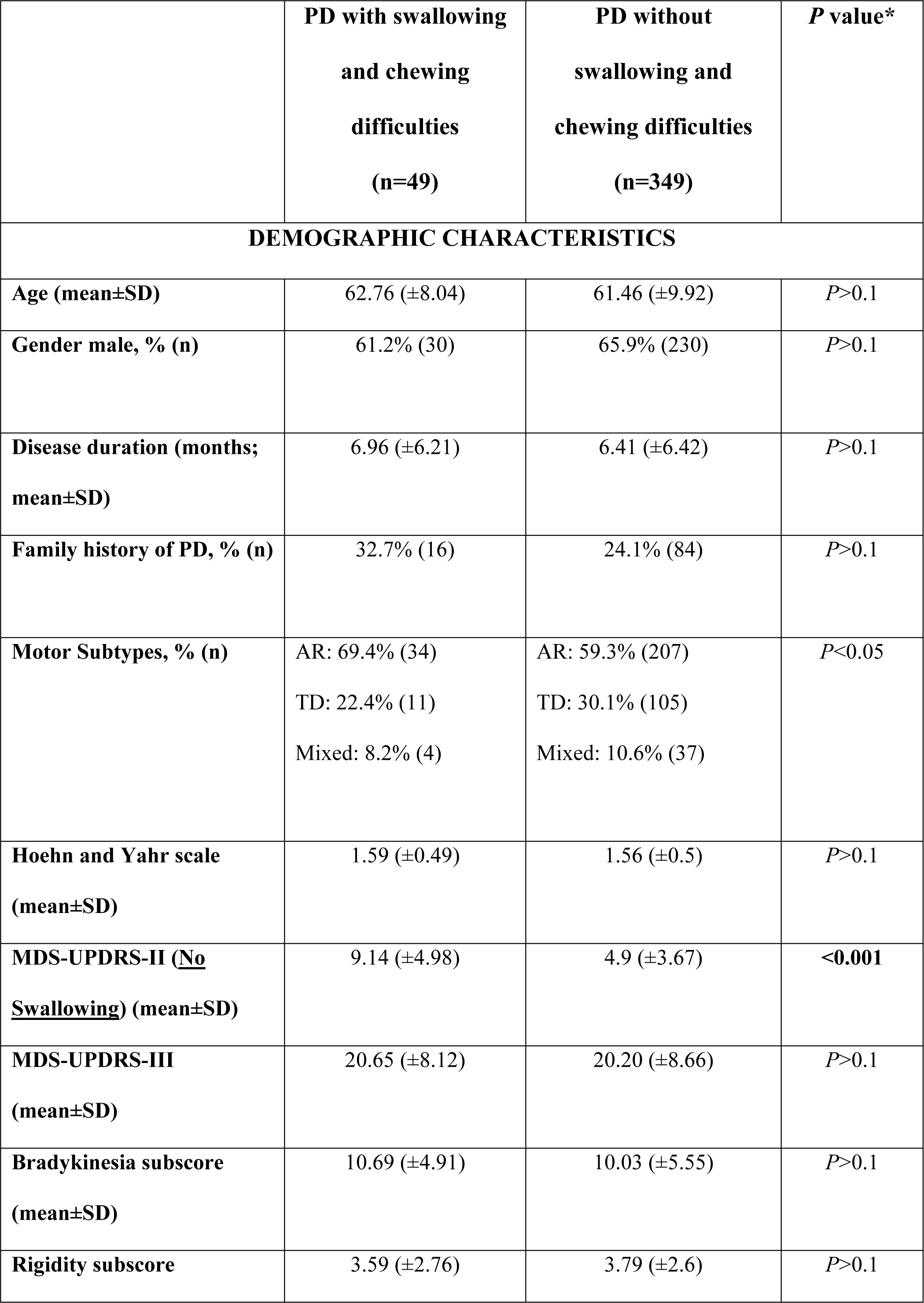
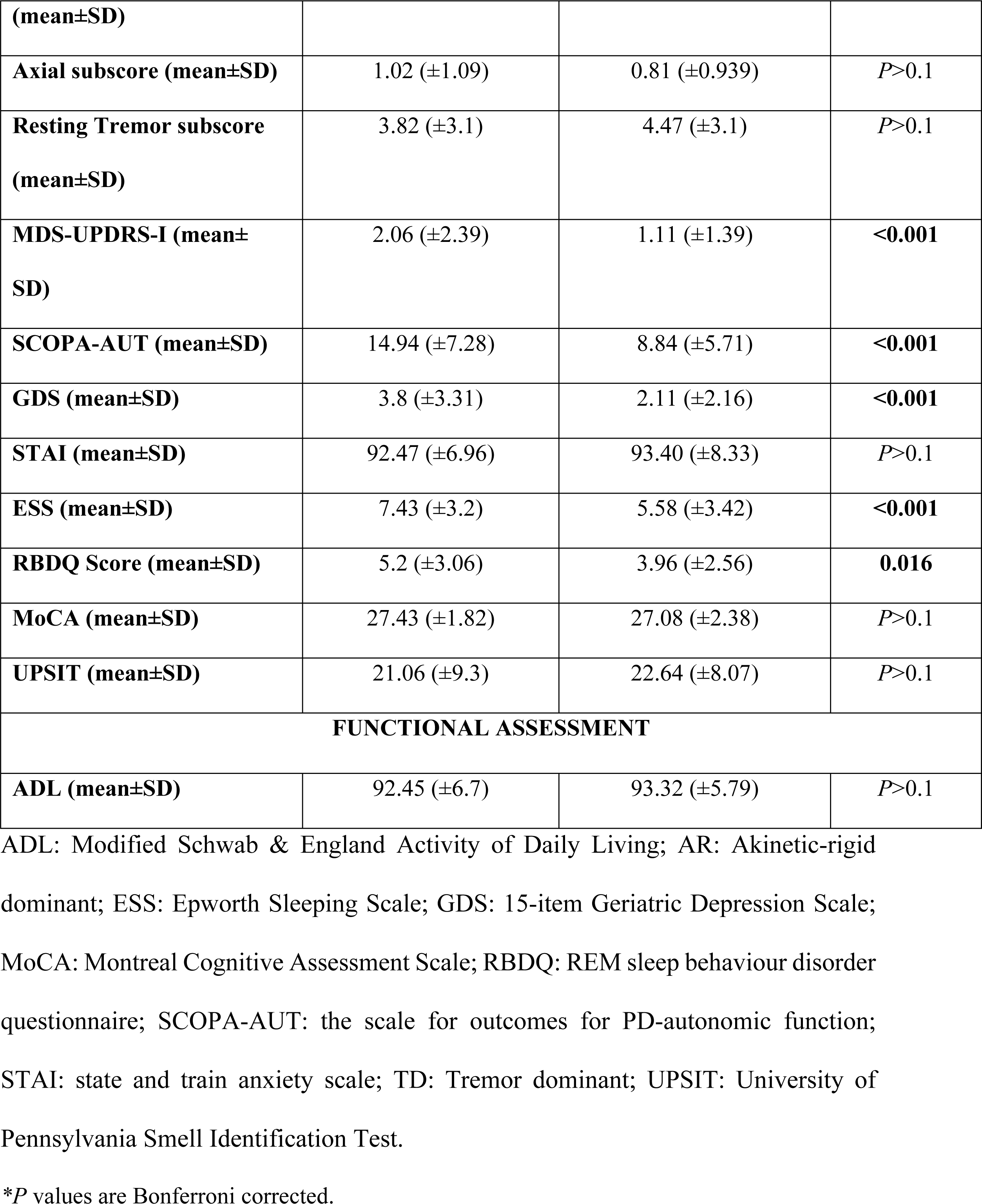
Demographic and clinical characteristics of early drug-naïve PD patients.

Early drug-naïve PD patients with swallowing and chewing difficulties had greater non-motor symptoms burden compared to those without swallowing and chewing difficulties. In specific, early drug-naïve PD patients with swallowing and chewing difficulties had higher MDS-UPDRS-I scores (*P*<0.001), worse autonomic dysfunction symptoms (SCOPA-AUT; *P*<0.001), depressive symptoms (GDS; *P*<0.001), excessive daytime sleepiness (ESS; *P*<0.001) and RBD (RBDQ; *P*=0.016) compared to PD patients without swallowing and chewing difficulties (Table 1). No significant differences were found in anxiety (STAI; *P*>0.1), cognitive function (MoCA; *P*>0.1), olfactory dysfunction (UPSIT; *P*>0.1) and disability scores (ADL; *P*=0.336) between early drug-naïve PD patients with and without speech and chewing difficulties.

### Imaging assessment: presynaptic dopaminergic function

Early drug-naïve PD patients with swallowing and chewing difficulties had significant lower [^123^I]FP-CIT uptakes in the striatum (*P*=0.016) compared to those without swallowing and chewing difficulties (Figure 1 and 2; Table 2). Within the striatum [^123^I]FP-CIT uptakes was significantly decreased in the caudate (*P*=0.008) but not putamen (*P*>0.1) of early drug-naïve PD patients with swallowing and chewing difficulties. Worse swallowing and chewing scores at the MDS-UPDRS-II item 2.3 were associated with lower [^123^I]FP-CIT uptakes in the striatum (*rho*=−0.157; *P*=0.002) and caudate (*rho*=−0.156; *P*=0.002: Figure 2).

**Figure 1.**
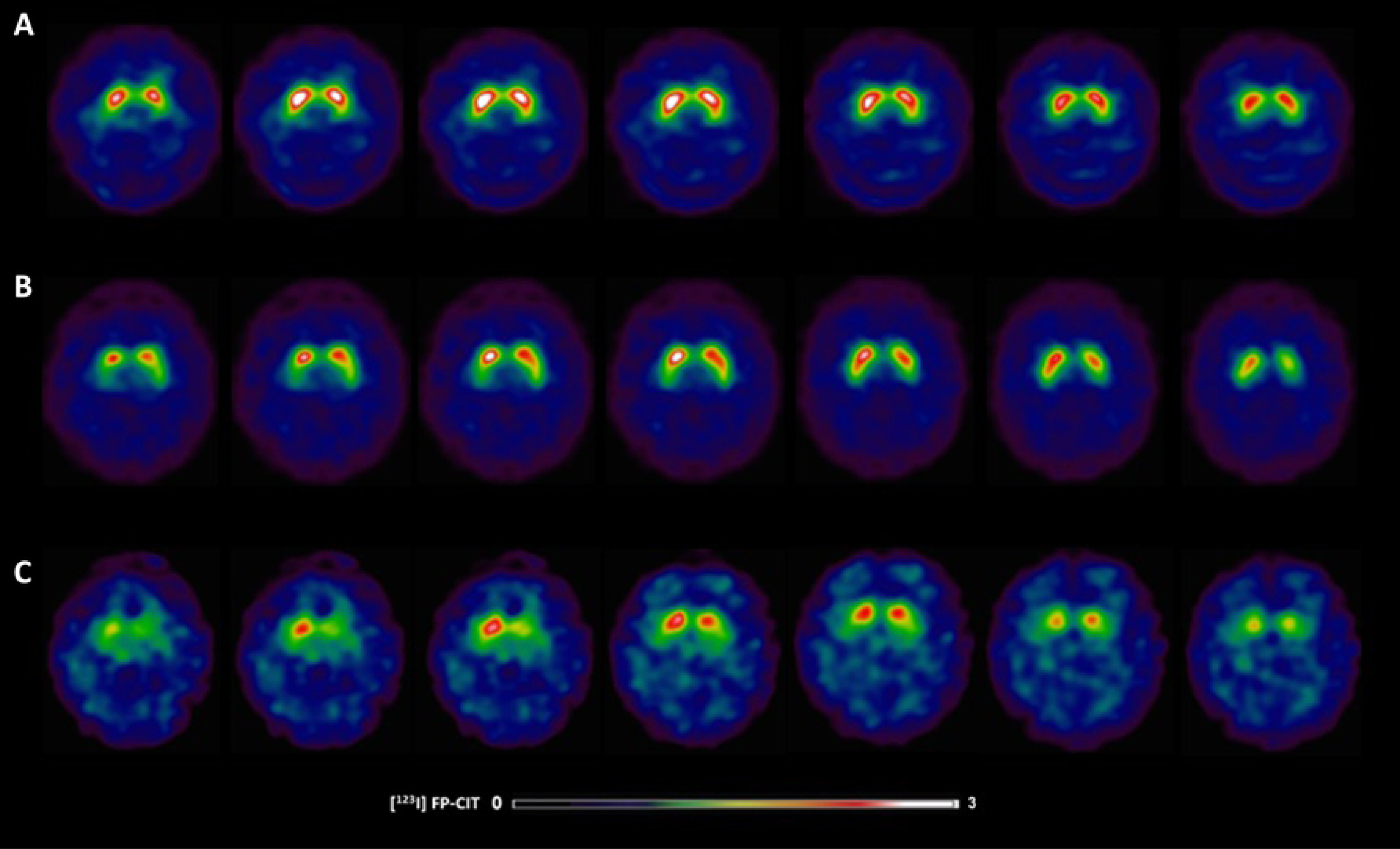
[^123^I]FP-CIT SPECT images in Parkinson’s disease patients with and without swallowing and chewing difficulties. (A) 63-year-old healthy control showing typical [^123^I]FP-CIT specific binding ratios in the caudate (SBR: 3.87) and putamen (SBR: 2.65) (B) 63-year-old male without swallowing difficulties exhibiting slight dopaminergic deficits as reflected by [^123^I]FP-CIT specific binding ratios in the caudate (SBR: 2.92) and putamen (SBR: 2.15); (C) 63-year-old female with swallowing difficulties demonstrating larger striatal dopaminergic deficits as reflected by [^123^I]FP-CIT specific binding ratios in the caudate (SBR: 1.00) and putamen (SBR: 0.58).

**Figure 2.**
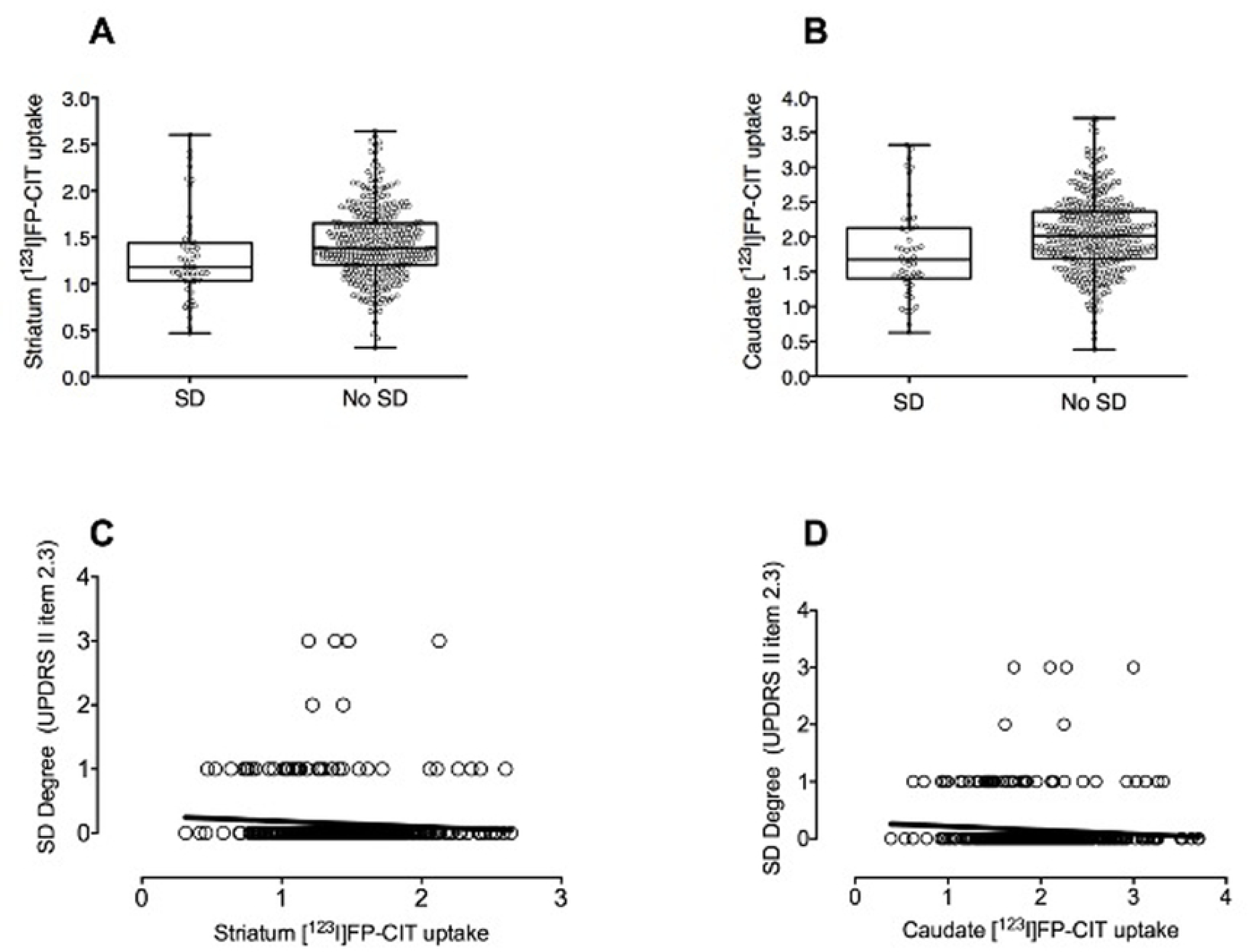
Presynaptic dopaminergic deficit in the group of early drug-naïve PD patients with and without swallowing and chewing difficulties. Box-plot showing decreased [^123^I]FP-CIT uptakes in the (A) striatum and (B) caudate of early drug-naïve PD patients with swallowing difficulties. Correlations between the degree of swallowing impairment (MDS-UPDRS-II, item 2.3) and [^123^I]FP-CIT uptake in the (C) striatum (*rho*=−0.157; *P*=0.002) and (D) caudate (*rho*=−0.156; P=0.002) of early drug-naïve PD patients. SD=Swallowing difficulties.

**Table 2.** [^123^I]FP-CIT uptakes in Striatum for the groups of early drug-naïve PD patients with and without swallowing difficulties.

| **Regions of Interest** | **PD with swallowing and**<br>**chewing difficulties**<br>**(n=49)** | **PD without**<br>**swallowing and**<br>**chewing difficulties**<br>**(n=349)** | ***P* value\*** |
| --- | --- | --- | --- |
| **Striatum (mean±SD)** | 1.28 (±0.48) | 1.43 (±0.38) | **0.016** |
| **Caudate (mean±SD)** | 1.79 (±0.66) | 2.03 (±0.53) | **0.008** |
| **Putamen (mean±SD)** | 0.78 (±0.37) | 0.83 (±0.28) | *P*>0.1 |

\**P* values are Bonferroni corrected.

### Motor progression and cognitive decline

Over a period of three years, 180/355 (50.7%) drug-naïve PD patients showed motor progression and 51/307 (16.6%) of them developed cognitive impairment. Cox proportional hazards analysis showed that the presence of swallowing and chewing difficulties in early drug-naïve PD patients has no influence on cognitive decline at the follow-up (Hazard Ratio (HR): 1.735, 95% Confidence Interval (CI): 0.890–3.381; *P*=0.106) or motor progression (HR: 1.259, 95% CI: 0.830–1.907; *P*=0.278).

## DISCUSSION

Our findings indicate that early drug-naïve PD patients with swallowing and chewing difficulties have greater presynaptic dopaminergic dysfunction in the caudate and non-motor symptoms burden. Moreover, the loss of striatal dopaminergic function is correlated with swallowing and chewing difficulties in early drug-naïve PD patients. Finally, swallowing and chewing difficulties are not associated with motor progression or an increased risk of cognitive dysfunction at the follow-up in early drug-naïve PD patients.

The prevalence of swallowing and chewing difficulties in the general PD patients’ population ranges between 9 to 77%.(16) We found that swallowing difficulties occur in 12.3% (49/398) of early drug-naïve PD patients and are more frequent in akinetic-rigid phenotype. This is the first study to report the prevalence of swallowing and chewing difficulties in early drug-naïve PD.

Early drug-naïve PD patients with swallowing and chewing difficulties had increased non-motor symptom burden suggesting a close association between swallowing abnormalities and PD non-motor features. Among the non-motor symptoms, early drug-naïve PD patients with swallowing and chewing difficulties showed worse autonomic dysfunction, depressive symptoms, excessive daytime sleepiness and disordered REM sleeping behaviour. On the contrary, anxiety, cognition, olfactory dysfunction and disability did not differ between the two groups. A recent study has underlined that patients with swallowing and chewing impairment are prone to affective symptomatology such as depression and fear. (18)

A recent study suggested that cognitive impairment (frontal/executive and learning/memory) is frequently corelated with the oral phase of swallowing in early PD patients. (19) However, we did not find any alterations in cognitive functions among early drug-naïve PD patients with and without swallowing difficulties. Moreover, swallowing and chewing dysfunction at baseline did not increase the possibilities of harbouring cognitive decline at the follow-up visits.

Furthermore, early drug-naïve patients with PD and swallowing and chewing impairment had substantially decreased striatal [^123^I]FP-CIT levels when compared to patients without swallowing impairment. The decrease of striatal presynaptic dopaminergic function was associated with the degree of swallowing impairment. To the best of our knowledge, this is the first time which striatal presynaptic dopamine deficits were associated with the degree of swallowing and chewing difficulties in PD. We found that early drug-naïve PD patients with swallowing and chewing deficits had the greater loss of dopaminergic nigrostriatal terminals in striatum and specifically in caudate.

The functional capacity of the supramedullary network controlling swallowing is depended on the in the integrity of dopaminergic neurons in the basal ganglia.(17) A bilateral activation of putamen and globus pallidus has been exhibited throughout swallowing procedures in a cohort of healthy controls.(18) Hence, in the context of dopaminergic deficits due to PD a dysfunction in the supramedullary swallowing system would be expected.

The spread of Lewy body pathology in PD, as described by Braak et all, involves several cortical and subcortical areas that elaborate in the regulation of swallowing mechanisms. (19) More specifically, the accumulation of Lewy bodies in areas of the medulla that control swallowing has been associated with severe dysphagia in PD patients. The ascending pattern of Lewy body pathology in PD initially involves initially dorsal nucleus IX and X and locus coeruleus (Stage I-II). These areas are predominantly involved in non-motor symptomatology. Subsequently, spreading of pathology encompasses substantia nigra, mesocortex and neocortex (Stage III-IV). Thus, motor features of PD become apparent. Following the ascending fashion of spreading, it would be expected that due to the early involvement of brainstem areas controlling swallowing, relevant symptoms would be evident in the early stages of the disease. Alas, severe swallowing difficulties are commonly present in advanced PD patients.

This inconsistency could be explained in the context of activation of compensatory mechanisms in cortical areas in early stages of PD. Using whole-head magnetoencephalography, it has been shown that cortical processing in PD patients without swallowing difficulties harbours a marked alteration of peak activity regarding lateral regions of premotor, motor, and inferolateral parietal cortex. Furthermore, the activity is activity is reduced in the supplementary motor cortex. Interestingly, PD patients with swallowing problems did not exhibit similar activity. These results are indicative of an adaptive mechanism with parallel motor networks, aiming to avoid swallowing dysfunction. However, when neurodegeneration exceeds a certain threshold, clinical symptoms of dysphagia might arise.

The absence of instrumental swallowing examination (i.e. Fibreoptic Endoscopic Evaluation of Swallowing and Videofluoroscopic Swallowing Study) as a validated way to assess swallowing impairment is a limitation of our study. Nevertheless, the use of the clinician-based scale such as the MDS-UPDRS-II (Chewing and Swallowing) item 2.3 may provide a simple tool for clinicians to assess the progression of swallowing difficulties. Finally, the limitation of our study is the correlational approach that does not allow us to perform direct attributions and conclude on causality. Therefore, our findings need to be interpreted with the appropriate caution.

In conclusion, our results indicate a significant correlation between swallowing difficulties and loss of striatal dopaminergic function in early drug-naïve PD patients. This specific subgroup of PD patients harbours a substantial burden of non-motor symptoms, without profound motor symptomatology.

